# The conserved wobble uridine tRNA thiolase Ctu1 is required to sustain development and differentiation

**DOI:** 10.1101/2023.11.08.566201

**Authors:** YZW Yu, CQ Wang, Y Wang, H Shi, ZL Zhou

## Abstract

Recent studies have revealed that tRNA modification is an important epigenetic mechanism involved in gene expression. Cytosolic thiouridylase (consisting of Ctu1 and Ctu2 subunit) are the enzyme complex which catalyze the thio-modification at the 34^th^ wobble uridine of the anticodon of tRNAGln^UUG^, tRNAGlu^UUC^, and tRNALys^UUU^. Besides introducing a thiol group at the C2 positions, those tRNAs were commonly modified with a methoxycarbonylmethyl at the C5 positions by Elongator and ALKBH8. tRNA-U34 modification, particularly the Elongator and ALKBH8, has been demonstrated to be involved in disease and development, however, the biological functional level of CTU in vertebrates remains elusive. Here, we found that in zebrafish, CTU may be an important regulatory factor in development and erythroid differentiation. By using morpholino targeting and knocking down CTU1, we observed that the loss of CTU1 led to impaired zebrafish larval development and blood vessel formation. Single-cell sequencing analysis showed that erythroid cell differentiation in the CTU1 knockdown group was blocked at an early stage, while the wild-type group exhibited mature erythroid cells. These findings suggest that CTU1 is involved in regulating erythrocyte development. These findings provide new insights into the biological function of CTU1.

## Introduction

Transfer RNAs (tRNAs) are known to undergo extensive post-transcriptional modifications steps for maturation[1, 2], especially the first position (position 34) of the tRNA anticodon which forms wobble base pairing with the third position of the codon. In eukaryotes, the wobble uridines (U34) of tRNAGln^UUG^, tRNAGlu^UUC^, and tRNALys^UUU^ are chemically modified with a thiol group at the C2 positions (s2), and (commonly) a methoxycarbonylmethyl at the C5 positions (mcm5) [3, 4]. The cytosolic tRNA thiouridylase complex, which are composed of two subunits, Ctu1 and Ctu2, is required for thiouridinylation of cytosolic tRNA. While the Elongator complex, consisiting of the six proteins Elp1-6, and ALKBH8 is required for the mcm5 modification. tRNA wobble U34 modification and their modifying enzymes have been conserved through evolution[5]. The complex mcm5s2U modifications enable tRNA to execute its delicate biological function and guarantees fidelity of protein translation[6–8].

tRNAs used to be constitutively expressed, and shows redundancy. However, recent studies have revealed that tRNA modifications can be dynamically altered in response to levels of cellular metabolites and environmental stresses. Thus, dynamic tRNA modifications represent another layer of epigenetic regulation of gene expression. Accumulating evidence shows that reprogramming of mRNA translation through altered tRNA activity can drive pathological processes in a codon-dependent manner. tRNA modifications and their modifying enzyme maybe involved in disease and development[9, 10]. For instance, ALKBH8 is highly expressed in human bladder cancers, and silencing of ALKBH8 significantly suppressed invasion, angiogenesis and growth of an in vivo model of human urothelial carcinoma cells [11]. The deletion of Ikbkap, a subunit of the elongation apparatus, leads to poor angiogenesis in mouse embryos, delayed entry into the mid-gastrula, disordered arrangement of dorsal primitive nerves, and failure of embryonic vascular system creation.[12]. Inactivation of either Ctu1 or Ctu2, has been shown to exhibit pleiotropic phenotypes, including cell cycle delay[13], defects in invasive growth[14], development abnormality[15], thermosensitivity[15], and sensitivity to various exogenous stresses like rapamycin[16], caffeine, oxidative stress or other stress[17–21]. In Arabidopsis thaliana, Ctu2 is essential for posttranscriptional thiolation of tRNAs and influences root development[22]. Besides, several clinical studies refer CTU2 as one of the candidate genes responsible for human dysmorphology syndromes. However, the in vivo role of Ctu1, and its control of genetic information in higher eukaryotes, especially in vertebrates, remains elusive.

Single-cell RNA sequencing (scRNA-seq) technology can provide comprehensive analysis of cellular and molecular information in a specific tissue. scRNA-seq were useful for the detection of the heterogeneous information that cannot be obtained by traditional sequencing[23, 24]. ScRNA-seq technology also plays an important role in revealing gene function, such as The knockout of Bcl11b will lead to the blockage of Puxi commitment of T cells. The gene regulatory network of cross-regulation of related transcription factors can be extracted by single-cell transcriptome analysis[25], SOX9-positive pituitary stem cells vary according to their location in the gland[26]. The deletion of runx2b significantly reduced the differentiation of osteoblasts and inhibited the formation of intermuscular bone[27]. Single cell sequencing confirmed that THBS2 was mainly expressed by fibroblasts, and the expression of THBS2 in fibroblasts from tumors was significantly higher than that of matched normal lung-derived fibroblasts. Pathway enrichment analysis showed that THBS2 fibroblasts had pleiotropic effects and immune inflammatory phenotypes. THBS2-rich TME may promote immune escape[28]. RNA velocity prediction showed that the differentiation of monocyte-dendritic cell progenitors into dendritic cells was blocked, and the rate of terminal neutrophil differentiation from preNeu1 to immNeu increased, and the change of cell trajectory caused by ThPOK deficiency[29]. Prdm6 regulates cardiac development by regulating the differentiation and migration of neural corona cells[30].

The zebrafish has emerged as an excellent model for studying vertebrate development due to their rapid external development and optical transparency[31]. Furthermore, zebrafish fli1 promoter is able to drive expression of enhanced green fluorescent protein (EGFP) in all blood vessels throughout embryogenesis[32, 33]. Herein, we analysed zebrafish ctu1 mutants with single-cell RNA sequencing. Here we applied morpholino antisense strategy to knockdown the epression of Ctu1 and test the role of Ctu1 during development in zebrafish. Meanwhile, scRNA-seq was applied to clarify the genes regulated by Ctu1 at different cell cluster’s level. For the first time, we demonstrate that Ctu1 is crucial in vertebrate development. Vasculogenesis and angiogenesis might be compromised without Ctu1. We also give our identification of genes regulated by Ctu1 by RNA sequencing at the single cell RNA level.

## Materials and methods

### Cell lines and culture conditions

Human microvascular endothelial cells (HMEC-1) were obtained from SUNNCELL, which were cultured in MCDB131 medium supplemented with 10 ng/mL Epidermal Growth Factor (Thermo, USA), 1 µg/mL Hydrocortisone (Sigma, Germany), 10 mM L-Glutamine (Gibco, New York, USA), 10% v/v Fetal Bovine Serum (Gibco, New York, USA), and 1%v/v Penicillin-Streptomycin Solution (Gibco, New York, USA). Cells were maintained at 37°C in a culture incubator (Thermo, USA) with 5% CO_2_.

### Lentiviral production and transduction

In this study, lentiviral vectors for gene modulation, focusing on CTU1 in HMEC-1 cells, were employed. Lentiviral constructs, pLenti-EF1a-EGFP-P2A-Puro-CTU1-KD for CTU1 knockdown and pLenti-EF1a-EGFP-P2A-Puro-CTU1-OE for overexpression, were validated (Obio Technology, Shanghai). Transduction occurred in 6-well plates at 2×10^5^ cells/ml.To ascertain the modulation of CTU1, quantitative PCR (qPCR) was conducted, employing well-established protocols for RNA extraction, cDNA synthesis, and the utilization of primers specifically designed for the quantification of CTU1.

### Quantitative real-time PCR

Total RNA was extracted from 1 x 10^6^ HMEC-1 cells subjected to CTU1 knockdown, overexpression, or control treatments using the RNA-easy Isolation Reagent Kit (Vazyme Biotech, China) following the manufacturer’s guidelines. Subsequently, 1 μg of the isolated RNA was reverse transcribed using the HiScript® III All-in-one RT SuperMix Perfect for qPCR kit (Vazyme Biotech, China). Quantitative PCR (qPCR) analysis was performed on a QuantStudio™ 5 Real-Time PCR System (Cell Signaling Technology, MA) using the SYBR green reaction mix (Vazyme Biotech, China, #Q711-02). The primer sequences: CTU1 sequence, synthesized by Sangon Biotech.

### Cell Viability Assay

For the assessment of cell viability following CTU1 overexpression and knockdown, 7000-8000 HMEC-1 cells were seeded into individual wells of 96-well plates. Cell growth and viability were monitored at 24-, 48-, and 72-hours post-seeding. All experiments were conducted in triplicate to ensure accuracy and reproducibility. Cell viability was evaluated using a Cell Counting Kit-8 (CCK-8, Beyotime, China) according to the manufacturer’s instructions. Optical density (OD) values were measured at 450nm with a microplate reader. To determine relative cell viability, the negative control group served as the reference point. Graphical representations were generated using GraphPad Prism 9.0, providing clear visualization of the observed trends in cell viability over time.

### Migration assay

Migration assays were meticulously conducted in 24-well plates (Corning, USA) utilizing Matrigel-coated membranes following the manufacturer’s recommended protocols. 1.5 x 10^4^ HMEC-1 cells which included CTU1 overexpression, CTU1 knockdown, and negative control (NC), starved for 12 hours, were diligently placed in the upper chambers of the inserts (Corning, USA). In the lower chambers, culture medium supplemented with 20% FBS was thoughtfully introduced to serve as a potent chemoattractant. Following a 24-hour incubation period at 37°C, meticulous care was taken to gently remove non-migrated cells residing on the upper membrane surface using cotton swabs. Subsequently, the remaining cells were fixed with 4% paraformaldehyde for precisely 15 minutes, followed by staining with 1% crystal violet for a duration of exactly 10 minutes at room temperature. Thorough washing with PBS three times ensued. Migrated cells were visualized and quantified through microscopic examination at a magnification of ×200 (Leica, Germany).

### Matrigel tube formation

The tube formation assay was conducted following a previously established protocol. In 96-well plates, a 50□µl layer of Matrigel (Corning, New York, USA) was evenly coated and left to gel at 37□°C for 30□min. HMEC-1 cells, reaching an 80% confluence in T25 culture flasks, were trypsinized and resuspended in growth medium at a concentration of 1.5 x 10^5^ cells/ml. Subsequently, 100μl of this cell suspension was gently added to each well. Afterward, the plates were placed in a 5% CO2 incubator at 37□°C for 2, 4, 19, and 24□h. Following incubation, the formed structures were observed and photographed under a microscope (Eclipse Ts2R; Nikon, Japan). For each condition, images were captured from a minimum of three random areas. The quantification of branch points, tube branch length, and total area covered was performed using ImageJ software (National Institutes of Health, USA) with the assistance of the "Angiogenesis Analyzer" plugin. Each experiment was repeated at least three times.

### Sample preparation for scRNA-seq

Single cell preparation, library construction, and sequencing were performed by Sinotech Genomics Co., Ltd. Shanghai, China. The protocols used for generating single cells are according to the paper before[34]. In brief, we collected around twenty zebrafish embryos (2 dpf) per group, and incubated them in the dissociation mix (480 μL 0.25% typsin, 20μL 100 mg/mL collagenase in Prepared DMEM-10% FBS) at 30□ for 5 minutes. During the incubation, the embryos were exposed with harsh pipettinging until they were fully homogenized without visible tissue. Then the digestion was stoped and the cell pellets were washed with PBS. Finally, the single cells in suspension were ready for the downstream procedures[34].

### Single-cell library preparation

The prepared single cells were loaded into the BD Rhapsody system (BD, San Jose, CA). The final cDNA library was generated from double strand full length cDNA by random priming amplification with the BD Rhapsody cDNA Kit (633773, BD Biosciences) and the BD Rhapsody Targeted mRNA & AbSeq Amplification Kit (633774, BD Biosciences). All the libraries were sequenced in a PE150 mode (Pair-End for 150bp read) in the X Ten instrument (Illumina, San Diego, CA).

### scRNA-seq data processing and quality control

The BD Rhapsody Whole Transcriptome Assay Analysis Pipeline was used. The FASTQ documents were filtered through multi-steps of quality inspection and sample labeling retrieval to generate a single cell expression profile matrix. The R software (version 4.2.2) and Seurat (version 4.3.0) were utilized for downstream clustering and visualization. We retained all the single cells which are high quality, including with unique feature counts above 200 and with less than 20% mitochondrial counts. After data quality control, we employed a global-scaling normalization by default method “LogMormalize”, scale factor is 10000. Principal component analysis (PCA) was then used to reduce dimensionality of the dataset to the top 50 PCs (Seurat ‘RunPCA’ function). Clustering was then performed using graph based clustering implemented by Seurat (‘FindClusters’ function). For visualization, Uniform Manifold Approximation and Projection (UMAP) coordinates were calculated using Seurat (‘RunUMAP’ function). The markers genes of each cluster were calculated by the FindAllMarkers function with the Wilcoxon Rank Sum Test under the following criteria: logfc.threshold = 0.25; min.pct = 0.25.

### CellCycle analysis

Cell cycle state was determined by the mean expression values of cell cycle marker genes. Each cell is assigned with a S-phase and G2/M phase score using Seurat’s ‘CellCycleScoring’ function.

### Differential genes analysis and gene enrichment analysis

In general, the differentially expressed genes (DEGs) between different models were determined by FindMarkers function with minimum percentage (min.pct) of 0.25 and log2 fold change threshold (log2FC.threshold) of 0.25. Genes were considered as significant by absolute log2FC was greater than 0.5 and their adjusted p-value was less than 0.05. Gene set enrichment analysis for DEGs were also performed using the R package clusterProfiler (v4.2.2). The gseKEGG and gseGO functions were used with OrgDb = "org.Dr.eg.db".

### Pseudo-time analysis

Monocle 2 (v2.22.0) was used to determine the pseudotime ordering of all cells. The top 100 DEGs of each clusters were used to order cells in the pseudotime trajectories. Dimensionality reduction was conducted using Discriminative Dimensionality Reduction with Trees (DDRTree). The dynamic expression of genes was visualized using the plot_genes_in_pseudotime function. The heatmap was visualized using the plot_pseudotime_heatmap function. The genes of each module were extracted based on the heatmap using the cutree function for GO and KEGG gene enrichment analysis.

### RNA velocity analysis

The balance of unspliced and spliced mRNA abundance is an indicator of the future state of mature mRNA abundance, and therefore an indicator of the future state of cells[35]. The direction and speed of each transition were evaluated based on the magnitude and direction of the individual cell speed arrows on the UMAP map. The spliced and unspliced reads of single-cell RNA sequencing data were calculated based on the bam file. The scv.tl.recovery _ dynamics function is used to recover kinetic information from single-cell RNA sequencing data, and the transition probability matrix in the dynamic system is estimated to infer the change of cell state over time. The RNA speed results were mapped to the UMAP image.

### Cell-cell communication analyses

Cell-cell communication networks were modeled based on the abundance of ligand-receptor pair transcripts using CellChat (version 1.6.1)[36]. To infer differences and similarities in the cell-cell communication network between ctu1 knockdown and WT cells, we subclustered mesoblastema cells from each condition. Cell groups of interest were merged, normalized, and used as input for CellChat. Zebrafish genes are converted into human genes through gene homology. We calculated the over-expressed genes and identified the significant ligand-receptor interactions using the identifyOverExpressedGenes (thresh.p = 0.05) and identifyOverExpressedInteractions functions, respectively. We utilized the ligand-receptor pairs human database provided by CellChat. The communication probabilities were calculated using the computeCommunProb function (raw.use=FALSE, nboot = 100, Hill function parameter kn = 0.5). The cellular communication network at the signaling pathway level was inferred using the computeCommunProbPathway function with default parameters (tresh = 0.05). The aggregated cell-cell communication networks were calculated using aggregateNet (threh = 0.05) with default parameters.

### GRN inference

To infer the gene regulatory network (GRN) between ctu1 knockdown and wild-type (WT) cells, we performed SCENIC using the pySCENIC functions[37]. After performing quality control and homologous gene conversion in Seurat, we exported the normalized data to a matrix, which was then converted into a loom file. Briefly, we utilized a pre-defined list of human transcription factors (TFs) (https://github.com/aertslab/pySCENIC/blob/master/resources/hs_hgnc_tfs.txt) and inferred regulatory interactions between them and their potential target genes using GRNBoost2. Then, we performed cisTarget motif enrichment using SCENIC’s RcisTarget and ranking databases (hg19-tss-centered-10kb-7species.mc9nr.genes_vs_motifs.rankings.feather). Last, the activity of the regulons was computed using SCENIC’s AUCell function. The activity of regulons in cells under different conditions is determined by converting Regulon Specificity Scores (RSS) to Z-scores, as described previously.

## Results

### Overview of the Phenotype and Single-cell Transcriptomic Analysis of Ctu1 Defective Zebrafish

CTU1 is a well-known conservative tRNA-U34 sulfur modification enzyme. In order to investigate the impact of Ctu1 on tRNA-U34 sulfur modification levels in zebrafish and to determine the biochemical function of CTU1, we used morpholino to specifically target knockdown CTU1. Additionally, we employed single-cell technology to identify cell types and perform functional analysis, as well as to elucidate molecular-level regulatory mechanisms. Through this approach,we can systematically, accurately, and comprehensively uncover the function and mechanism of action of CTU1 (Fig. 1A). Firstly, we examined the expression level of CTU1 in normal samples and found that CTU1 is most highly expressed in endothelial cells (GSE186423, Fig. 1B). According to the transparent nature of zebrafish blood vessels, we selected zebrafish as the model organism for our analysis. We used the antisense morpholino oligonucleotides (MO) strategy[33] in transgenic zebrafish Tg(fli1a:EGFP) y1 to rapidly screen the potential role of Ctu1 in vertebrate animals. Antisense ctu1-i2e3-MO targets the splice acceptor site of exon 3 (intron 2-exon 3 junction) of ctu1, thereby interrupting the normal splicing of its primary transcript (Fig. 1C upper). Fertilized one-cell stage zebrafish embryos were injected with control or antisense ctu1-i2e3-MO. Two days after the morpholino injection, nucleic acids were extracted from the embryos for an RT-PCR experiment. The PCR result showed that the antisense ctu1-i2e3-MO effectively knocked down ctu1 expression (Fig. 1C, lower). Two days after microinjection, the zebrafish larvae were anesthetized and dechorionated. Photographs were then taken under a stereomicroscope, capturing both brightfield and fluorescence images. The images depict the embryonic development and vascular morphology of zebrafish larvae (Fig. 1D). Control or antisense ctu1-i2e3-MO-injected larvae were anesthetized and imaged at 2 days post-fertilization (dpf). The control morphants had clearly defined and well-shaped structures (Fig 1E, F upper), while ctu1 morphants experienced a greater incidence of developmental abnormalities with morphological malformations such as expanded brain ventricles and severe hindbrain edema, pericardial edema, yolk extension deformities, misshapen spine, and a cocked-up tail (Fig 1E, F lower). Furthermore, Tg(fli1a:EGFP)y1 zebrafish, which exhibit eGFP fluorescence in endothelial cells, were also used for continuous in vivo observation of vascular structures during embryonic development[32]. GFP imaging showed well-developed intersegmental vessels (ISV) and dorsal longitudinal anastomotic vessels (DLAV), as well as a honeycomb-like caudal vein plexus (CVP) in control larvae (Fig 1G, left). In contrast, Ctu1 morphant larvae exhibited significant vasculogenesis defects, including impaired trunk angiogenesis and CVP formation (Fig 1G, right). Circulation in the caudal vein is visible in the control group, but is rarely visible with intermittent mobility in larvae deficient in Ctu1 (Supplementary Movie 1). So, those embryos deficient in ctu1 showed remarkable delayed development in vivo. The zebrafish knocked down by Ctu1 showed defects in embryonic development, particularly affecting blood vessel formation.

**Figure 1.**
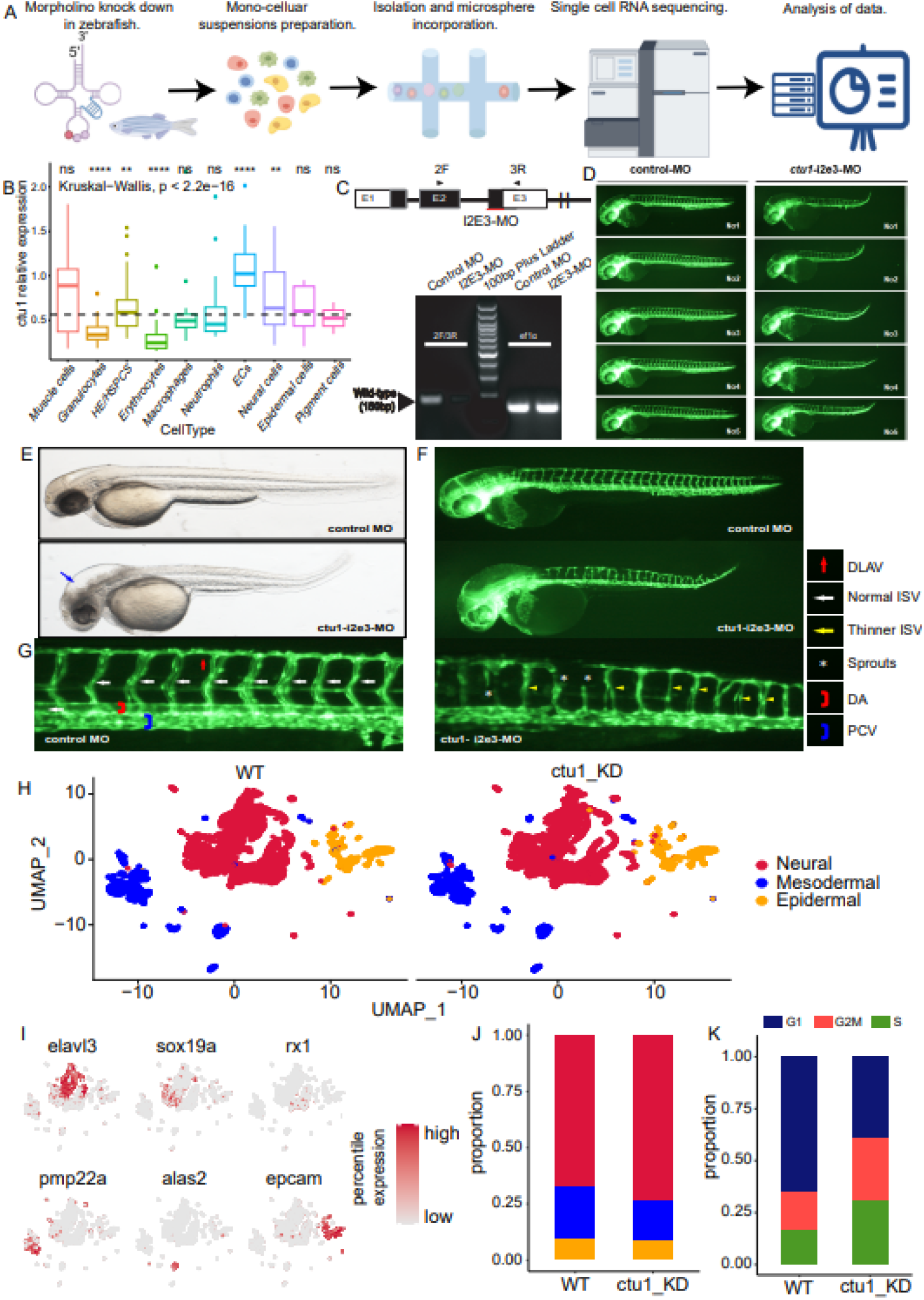
Morpholino knockdown of Ctu1 results in developmental defects in zebrafish embryos. (A)Schematic representation of scRNA-seq and analysis flowchart by Figdraw. (B)Box plot shows ctu1 relative expression in each celltype. ****p < 0.0001. (C)The zebrafish ctu1 gene was targeted by specific morpholino antisense strategies to block the splice acceptor site of exon3 (I2E3-MO). RT-PCR of ctu1 transcript from control-MO and I2E3-MO morpholino-injected embryos 2 days after fertilization, demonstrating knock-down of the splice acceptor site of exon3. (D) Each column shows five examples of the phenotype induced in the trunk angiogenesis. (E-G) Representative bright field and fluorescent images of Tg(fli1a:EGFP)y1 embryos at 2-dpf. Compared with control MO, knock down ctu1 present expanded brain ventricle and hindbrain oedema (blue arrow). (F,upper;G left) Image of trunk regions taken at 2-dpf, with the vascular structures visualised by eGFP fluorescence and labelled ISV (intersegmental vessel) and DLAV (dorsal longitudinal anastomotic vessel) showed regular development in the embryo injected with control MO. Compared with control MO, embryos injected with ctu1-i2e3-MO present a lower number of incomplete and thinner ISVs (yellow arrow) and ectopic sprouts (asterisk) of dorsal aorta (F,lower;G right). The boxed regions are shown at higher magnification in the bottom panels. (H) UMAP representation of the scRNA-seq matrix from ctu1 defects cells and WT cells. (I)UMAP visualisation of key marker gene expression. Colour scale represents log-normalised expression. Neural markers:elavl3,sox19a,rx1. Mesodermal markers:pmp22a,alas2.Epidermal marker:epcam. (J) Percentages represent the proportion of mutant cells and 3WT cells per main celltype. (K) Percentages represent the proportion of mutant cells and 3WT cells per cellcycle phase.

We took 2-day-old zebrafish larvae after MO microinjection and fertilization. Approximately 20 individuals were mixed in the control group and Ctu1-KD group, respectively, for tissue lysis and scRNA-seq (Fig. 1A). Data derived from 20,788 cells was retained for downstream bioinformatics analysis. After clustering the transcriptomes, we generated 32 distinct cell clusters and visualized them using the uniform manifold approximation and projection (UMAP) approach. Subsequently, we identified 22 cell types based on known marker genes[38], which roughly include neural, mesodermal, and epidermal cells (Fig. 1H, I; Fig. SI.1A-C). From the perspective of the cell ratio between the two samples, the proportion of neural cells increased in the ctu1 knockdown group, while the proportion of mesodermal cells decreased (Fig. 1J). Additionally, the ctu1 knockdown group showed an increase in the proportion of cells in the G2M phase and S phase when comparing the cell cycle phases between the two samples (Fig. 1K). These data demonstrate that the cells were in an active division phase after the knockdown of CTU1.

### Heterogeneity of Ctu1 defects in wild zebrafish cells

In order to understand the heterogeneity between the two samples, we calculated the differentially expressed genes and performed pathway enrichment analysis. The upregulated genes captured predicted cell cycle phase transition markers, including ccng1, ccnd1, and tp53. The Significantly downregulated differential genes including ATPase Na+/K+ transporting subunit alpha 3a (atp1a3a), synuclein beta (sncb), EBF transcription factor 3a (ebf3a), pre-B-cell leukemia homeobox 3b (pbx3b), keratin 4 (krt4), solute carrier family 1 member 2b (slc1a2b), type I cytokeratin (cyt1l), ELAV like neuron-specific RNA binding protein 4 (elavl4), ribosomal protein (rplp2), creatine kinase (ckbb)(Fig. 2A, SupplementalTable S1). The upregulated genes in ctu1 knockdown cells were enriched in double−strand break repair, DNA replication, DNA repair, DNA metabolic process, cell cycle, downregulated genes were enriched in transcription cis-regulatory region binding, RNA polymerase II transcription regulatory region sequence-specific DNA binding, membrane, axon, structural molecule activity (Fig. 2B, SupplementalTable S2).We isolate the three germ layers and analyze them. The neural cells were segregated and reclustered, including Neural_sox19a, Neural_neuron_elavl4, Neural_neuron_ebf2, Neural_neuron_phox2a, Neural_neuron_dlx1a, Neural_neuron_eomesa, Neural_optic, Neural_neural crest, Neural_retina, and Neural_floorplate (Fig SI1B). We performed a comparison between CTU1 knockdown and wildtype samples, identified 104 significantly downregulated genes and 69 significantly upregulated genes. The activation of DNA replication, DNA repair, cellular response to DNA damage stimulus, DNA metabolic process, and cell cycle are consistent with the overall difference. However, the distal axon, growth cone, and site of polarized growth are inhibited, indicating that CTU1 knockdown has an effect on neural development(Fig. 2C, SupplementalTable S3). We identified only 4 celltypes that corresponds to the Epidermal lineages which including Epidermal_icn2, Epidermal_cldn, Epidermal_otic placode, Epidermal_lens(Fig SI1C). Differential gene expression analysis identified 395 variable genes, including 56 significantly downregulated genes and 24 significantly upregulated genes. As GSEA analysis revealed activation of RNA splicing, nuclear protein−containing complex, mRNA metabolic process, polymeric cytoskeletal fiber, chromatin, chromosome whereas polymeric cytoskeletal fiber, supramolecular polymer, supramolecular fiber, structural molecule activity, supramolecular complex were decreased(Fig. 2D, SupplementalTable S4).

**Figure 2.**
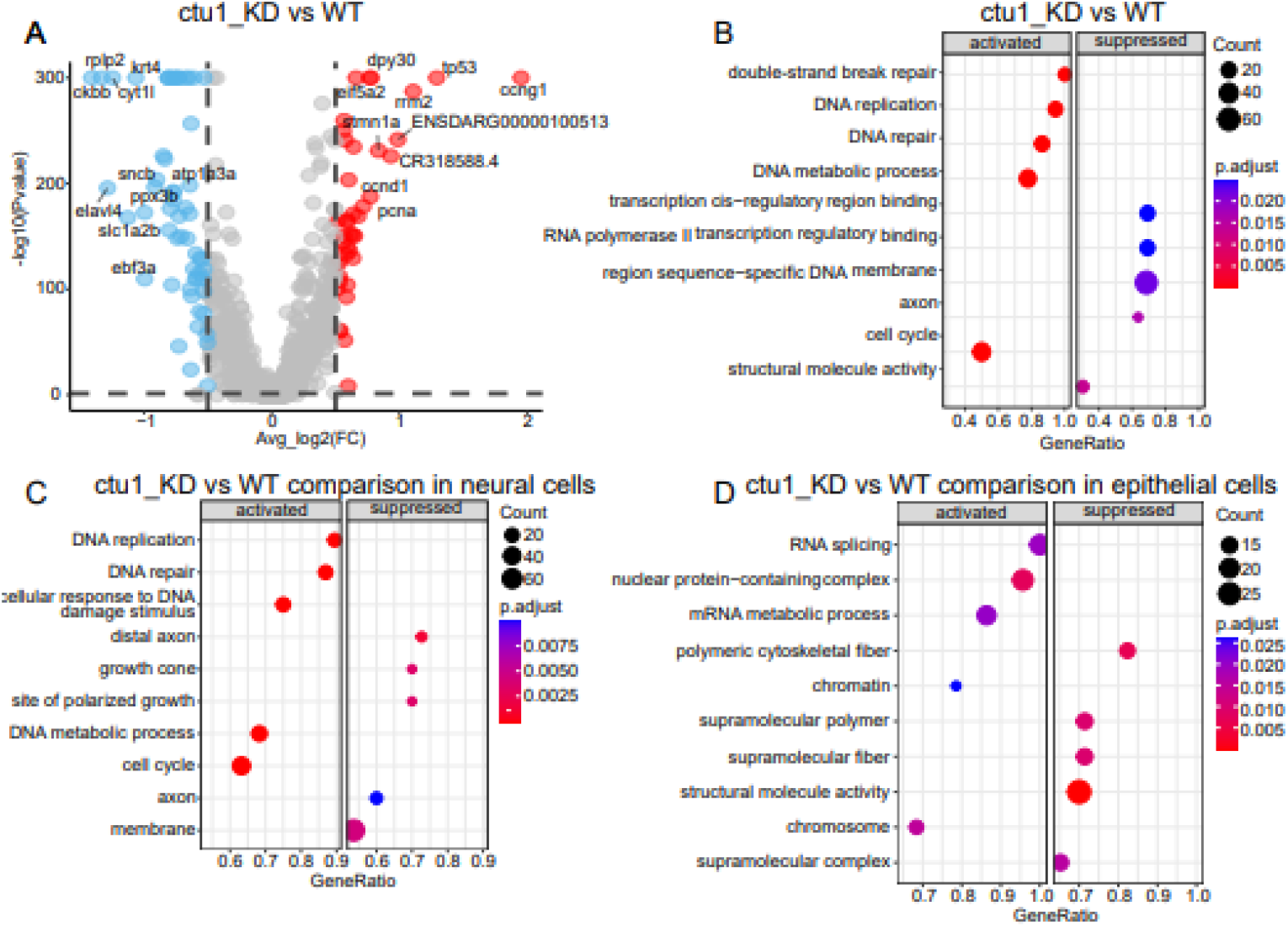
Heterogeneity of Ctu1 defects in wild zebrafish cells. (A) A volcano plot is shown, highlighting the top 10 differentially expressed genes (DEGs) that are upregulated and downregulated. (B) The major GSEA GO term enriched in each sample. (C) The major GSEA GO term enriched in neural cells of each sample. (D) The major GSEA GO term enriched in epithelial cells of each samples.

### Heterogeneity of Ctu1 defects in mesodermal cells of wild zebrafish

We next explored the heterogeneity of mesoblastema cell expression. Mesoderm cells were extracted, and UMAP dimensionality reduction clustering analysis was performed (Fig. 3A). To infer cell trajectories, we performed RNA velocity analysis[35], which confirmed that the muscle differentiation trajectory in the transitional population developed largely towards the pharyngeal arch or heart cluster. However, the differentiation rate slowed down after knocking down ctu1 (Fig. 3B). It was speculated that the wild-type samples were in the late stage of differentiation during scRNA-seq sampling, while the mutant group was in the early stage of differentiation.

**Figure 3.**
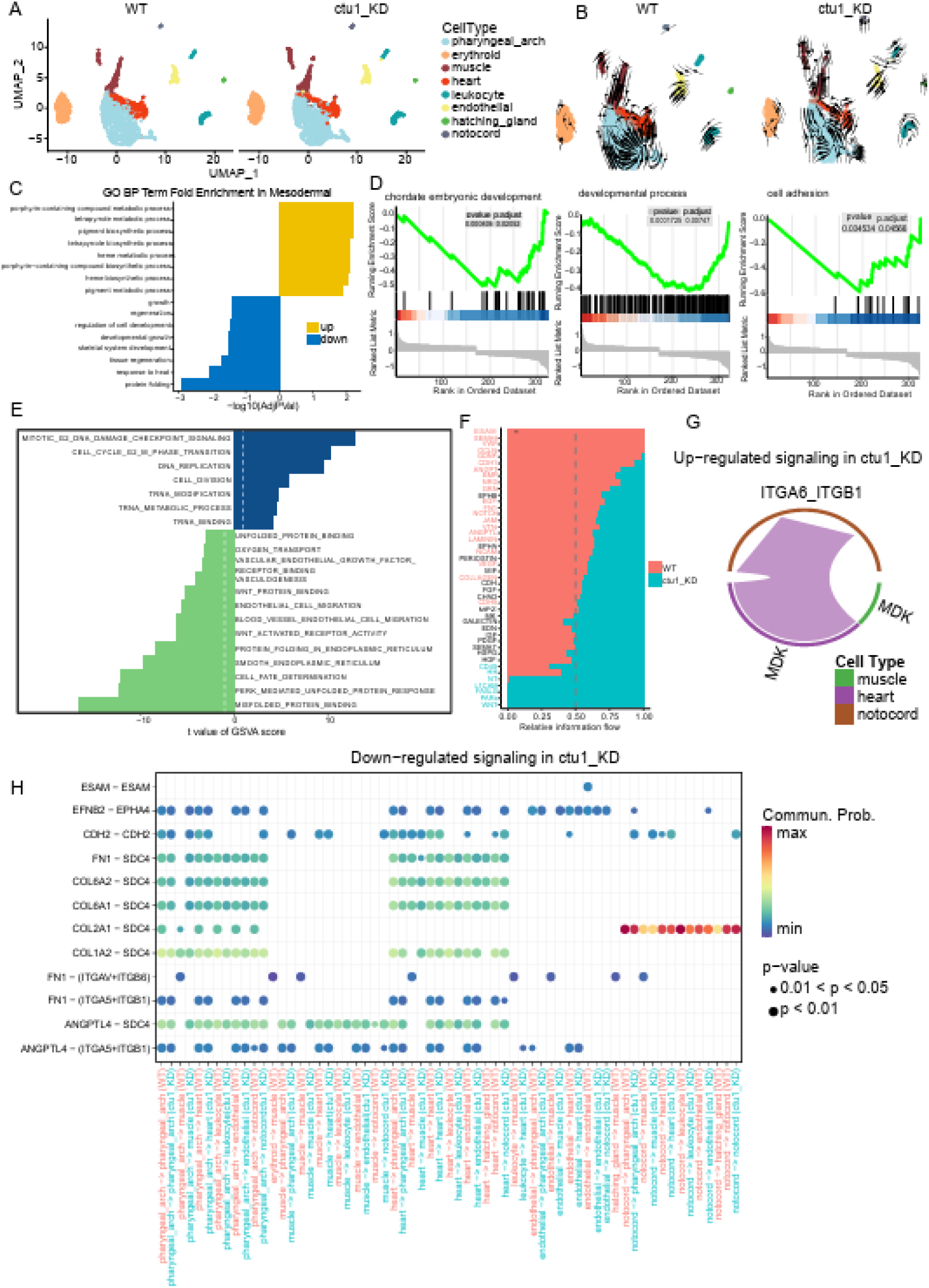
Heterogeneity of Ctu1 defects in mesodermal cells of wild zebrafish. (A) UMAP visualization of 4240 cells for mesoderm populations, colored according to cell type. (B) RNA velocity plot of WT mesodermal cells (a, left) and ctu1_KD mesodermal cells (b, right). Overlaid arrows indicate the predicted RNA velocities derived from spliced versus unspliced scRNA-seq reads. (C) The bar plot shows the Gene Ontology Biological Process (GO BP) analyses of differentially expressed genes (DEGs) between WT and ctu1−/− mesodermal cells. The y-axis represents the enriched BP term, while the x-axis represents the −log10 (adjusted P-value). GO terms enriched in ctu1-/- are colored yellow, and terms enriched in WT are colored blue. (D) GSEA plots show enrichment of Gene Ontology (GO) gene sets in differentially expressed genes (DEGs) between WT and Ctu1-deficient mesodermal cells. Each bar represents a GO term. (E) GSVA plots show enrichment of gene ontology (GO) gene sets between WT and Ctu1-deficient mesodermal cells. (F) All significant signaling pathways were ranked based on the differences in overall information flow within the inferred networks between WT and Ctu1-deficient mesodermal cells. The top signaling pathways, colored red, are more enriched in WT. The middle pathways, colored black, are equally enriched in WT and Ctu1-deficient cells. The bottom pathways, colored blue, are more enriched in Ctu1-deficient cells. (G) The chord plot shows upregulated signaling in Ctu1-deficient cells. (H) The dot plot shows downregulated signaling in Ctu1-deficient cells.

To understand the molecular basis for the altered cellular trajectory due to CTU1 deficiency, we examined the differential gene expression in ctu1 knockdown cells compared to WT progenitor populations using FindMarkers function. This analysis revealed that upregulated expressed genes tend to coincide with heme metabolic process, heme biosynthetic process, and porphyrin-containing compound metabolic process (urod, hmbsa), while downregulated DEGs tend to impact broad steps in development, which include developmental growth (klf6a, igf2b, junbb, hapln1a), regulation of cell development (her9, her6, hey1), skeletal system development (sox5, col2a1b, rpl13, hapln1a), or protein folding (hsp70.3, hspa5, hsp70.2, hsp70l) (Fig. 3C, SupplementalTable S5). As gene set enrichment analysis (GSEA) revealed, mRNA processing was activated, while cell adhesion, developmental process, chordate embryonic development, and translation were decreased (Fig. 3D, SupplementalTable S6). Gene set variation analysis (GSVA) further indicated that DEGs of ctu1 knockdown correlated with cell cycle G2/M phase transition, tRNA modification, tRNA metabolic process, and tRNA binding. In contrast, the binding of vascular endothelial growth factor receptor, endothelial cell migration, vasculogenesis, and cell fate determination were suppressed (Fig. 3E). Together, these findings provide evidence that the absence of ctu1 promotes cell proliferation and inhibits cell differentiation, thereby affecting angiogenesis and individual development.

In order to gain a deeper understanding of the communication between mesoderm cells after knocking down ctu1, we utilized the cellChat algorithm to infer the ligand binding of different cell interactions. First, we examine the quantity and intensity of cellular communication from a holistic perspective. It can be seen that, compared with the wild-type cell sample, the total communication volume increased in the ctu1 knockdown sample, but the overall communication intensity decreased significantly (Fig. SI2A).

Although most of the pathways showed signal activity in WT and ctu1 _ KD samples, the intensity of ESAM, SEMA6, VWF, NOTCH, VEGF and NCAM pathways was significantly reduced compared with the control group. Study showed VWF encodes a glycoprotein involved in hemostasis. The encoded preproprotein is proteolytically processed following assembly into large multimeric complexes. These complexes function in the adhesion of platelets to sites of vascular injury and the transport of various proteins in the blood. Mutations in this gene result in von Willebrand disease, an inherited bleeding disorder[39, 40]. Additionally, the first three pathways were specific communication pathways in wild-type (WT) samples. The intensity of CD45, HH, NT, L1 CAM, FASLG, PARs, and WNT pathways was significantly higher than that in the control group. These pathways were specific communication pathways in the mutant samples (Fig. 3F). We further identified the differentially expressed signal genes related to wild-type (WT) and ctu1 knockdown samples (Fig. 3G). After knockdown of ctu1, the MDK− (ITGA6+ITGB1) receptor-ligand pair increased the probability of communication between the heart, notochord, and muscle. Midkine(MDK) is a heparin-binding growth factor that promotes the growth, survival, migration and differentiation of various target cells[41]. And they found that alpha6beta1-integrins are functional receptors for midkine. Hoshino’s team reported that exosomal integrins may serve as predictive markers for organ-specific metastasis. Notably, integrins α6β1 were specifically associated with lung metastasis[42]. Additionally, the autocrine signal ESAM from endothelial cells decreased. The EFNB2-EPHA4, which plays an important role in cell adhesion and vascular development, decreases the probability of communication between the endothelium, gill arch, and heart. EFNB2 gene encodes ephrin (Eph) family, binding to receptor tyrosine kinase including EPHA4, plays a central role in heart morphogenesis and angiogenesis through regulation of cell adhesion and cell migration Cell surface transmembrane ligand for Eph receptors, a family of receptor tyrosine kinases which are crucial for migration, repulsion and adhesion during neuronal, vascular and epithelial development. The COL1A1/COL1A2/COL6A1/COL6A2-SDC4 receptor-ligand pairs mediate cell adhesion and signal transduction, thereby reducing the likelihood of communication between the heart, gill arch, and white blood cells, while specifically targeting white blood cells.

### Heterogeneity of Ctu1 defects in wild zebrafish endothelial cells

Zebrafish endothelial cells were extracted for research (Figure 4A), and differential gene analysis was performed. The gene expression levels of the ctu1_KD group were lower than that of the WT group. Significantly different genes are all ribosome genes, including rplp2, hsp70l, hsp70.1, rpl13, rplp1, hsp70.3, rpl24 (Figure 4B, SupplementalTable S7). Combining gene ontology analysis, differentially expressed genes were enriched in ribosome-related pathways including ribosomal subunit, cytosolic ribosome, as well as in response to unfolded protein, response to heat, myeloid cell homeostasis, erythrocyte homeostasis, erythrocyte differentiation, embryonic skeletal system morphogenesis pathways (Figure 4C, SupplementalTable S8). To identify specific gene regulators (regulons) that may be undergo expression changes following a decrease in ctu1 expression, we performed gene regulatory network (GRN) analysis using pySCENIC[37], and identified significantly active regulons specific for each cell type in the mesoderm were identified, endothelial cell-specific regulatory factors include FEV, SMAD1, GATA2, HLX, and PRDM1 (Fig SI.2E). Compared with the widetype group, TP53, MLX, STAT3, RELA are highly active in after knockdown ctu1. However, MEF2D,TGIF1,SIX1,HES1,SOX10 transcription activity decreased (Figure 4D).

**Figure 4.**
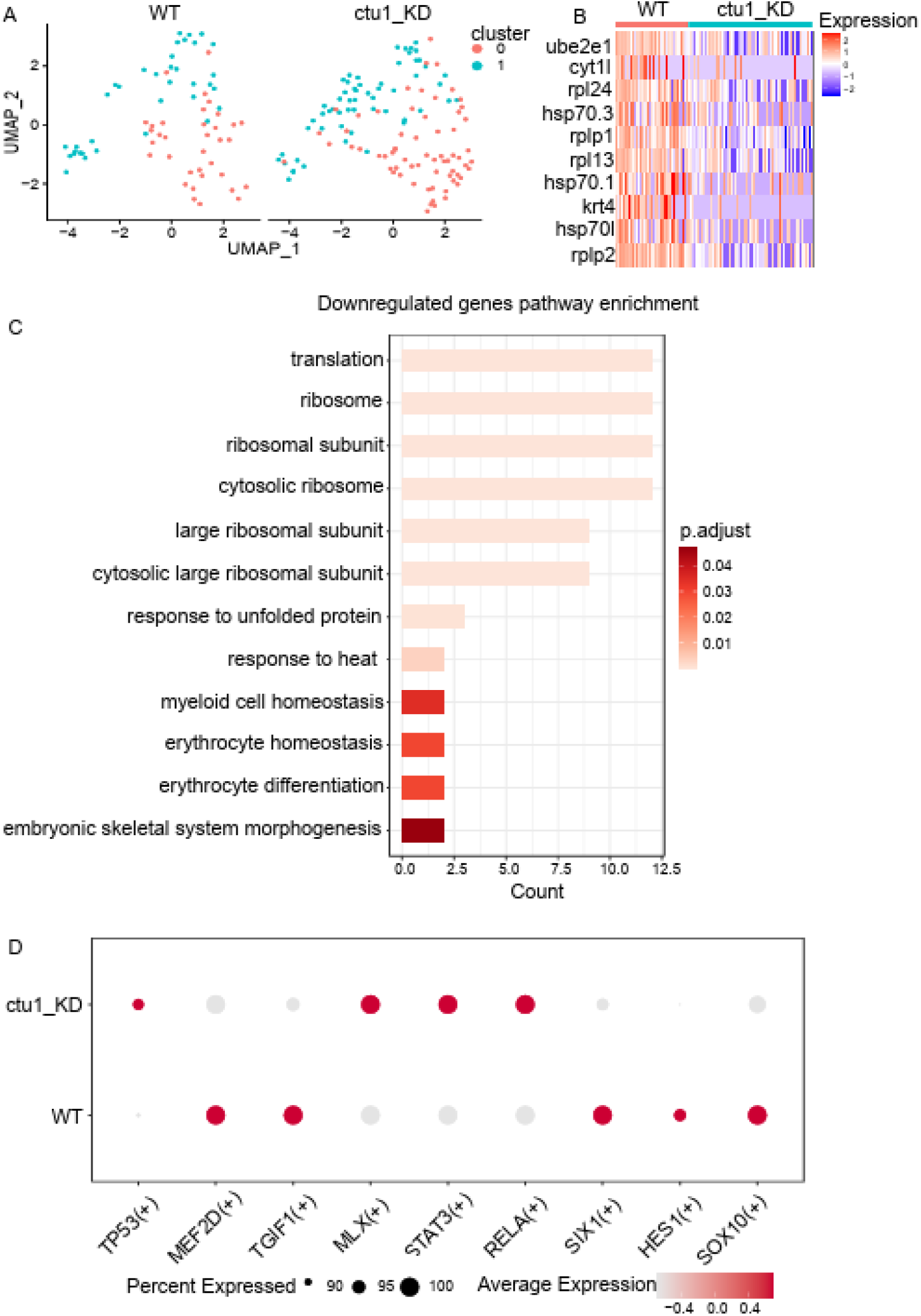
Heterogeneity of Ctu1 defects in wild zebrafish endothelial cells. (A) UMAP visualization of endothelial cells, colored according to clusters. (B) Heatmap plot of the top 10 DEGs between WT and ctu1_KD endothelial cells. (C) The bar plot shows the enrichment of GO BP terms in downregulated genes between WT and ctu1_KD mesodermal cells. (D) Dot plots show changes in the expression of transcription factors across different samples. The color and size of circles indicate the average expression level and percentage of cells.

### Knockout of CTU1 affects angiogenesis

To investigate CTU1’s impact on angiogenesis, we manipulated CTU1 expression in HMEC-1 cells using transfection (Fig. 5A). This allowed us to validate changes in RNA expression, revealing a significant five-fold increase in knockdown efficiency and a four-fold increase in overexpression efficiency compared to the NC group (p < 0.0001). In assessing the impact of CTU1 gene expression on cell viability, we cultured cells with targeted CTU1 gene alterations and monitored cell vitality at different time points. Our findings revealed that overexpression of CTU1 significantly enhanced cell activity after 48 hours, whereas CTU1 knockdown markedly reduced cell viability by more than one-fold (p < 0.0001) (Fig.5B). In addition, the exogenous expression of CTU1 significantly augmented the migratory capacity of HMEC-1 cells, while the downregulation of CTU1 expression moderately attenuated their migratory potential (Fig.5C).To explore the migratory behavior of HMEC-1 cells under varying CTU1 expression conditions, our experiments distinctly illustrated that HMEC-1 cells overexpressing CTU1 exhibited capillary-like structures as early as 4 hours, in stark contrast to the control group, which displayed minimal changes. Notably, at the 16 hour time point when cells were at their optimal growth state, there were subtle differences in tube branch length compared to the control group, but substantial variations in tube branch length and total area covered. Crucially, CTU1 knockdown cells displayed even more pronounced differences compared to the overexpression group. Throughout the entire culture period, CTU1 knockdown HMEC-1 cells failed to form tubular structures, indicating a more pronounced inhibitory effect on angiogenesis (Fig. 5D).

**Figure 5.**
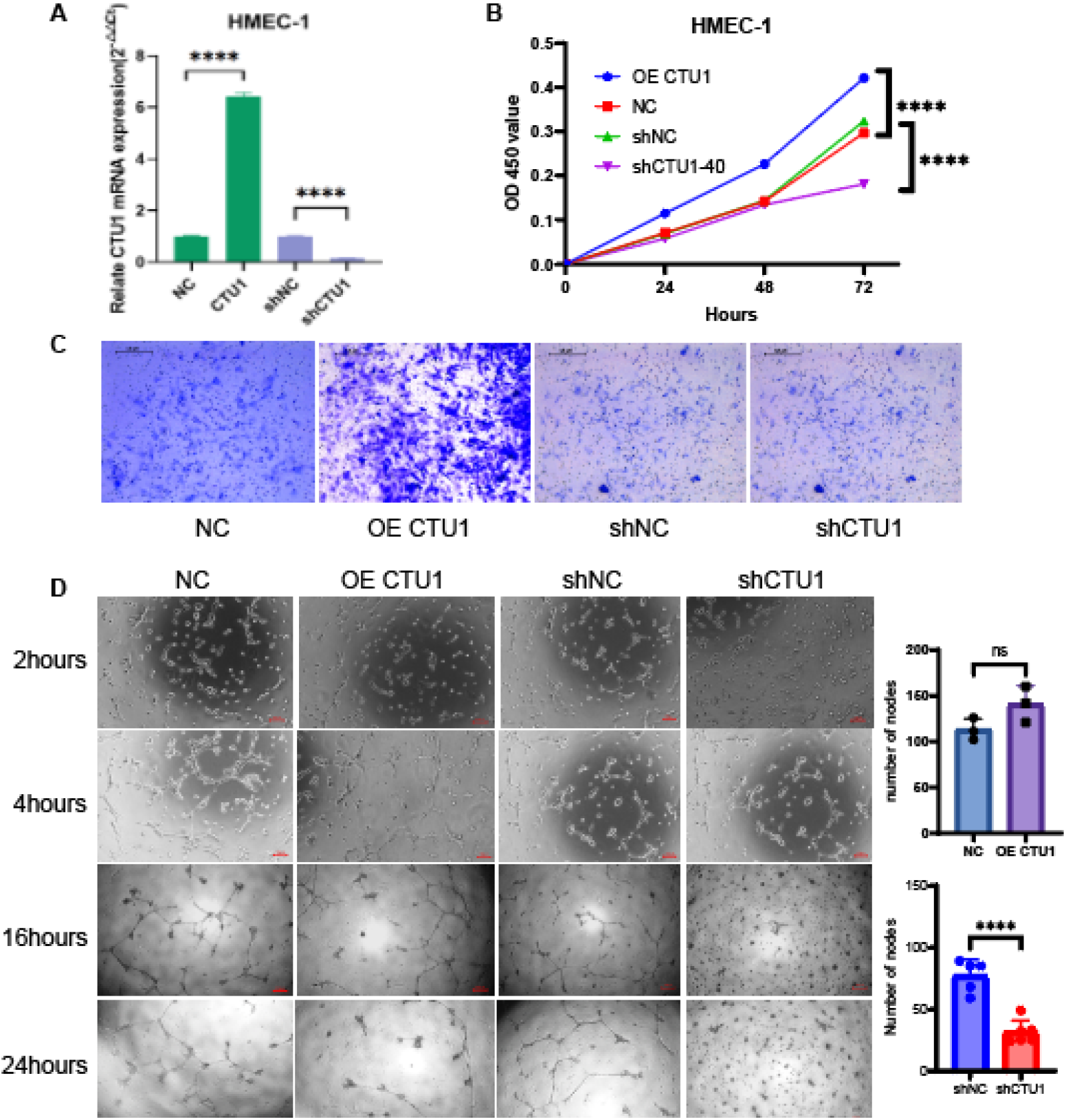
Experimental validation of CTU1 knockout or overexpression on HMEC-1 cell impact. (A) RNA expression levels of CTU1 gene in HMEC-1 cells with knockdown or overexpression. (B). Effects of CTU1 overexpression or knockdown on HMEC-1 cell viability. (C). Representative microscopic views of cell staining of HMEC-1 cells with CTU1 gene knockdown or overexpression that migrated through the transwell membrane compared to the control group. (D). Left: Representative images of tube formation experiment in HMEC-1 cells with CTU1 overexpression or knockdown at 2h, 4h, 16h, and 24h time points. Right: Statistical analysis of different cell groups at the 16-hour time point. Three independent experiments were conducted with similar results. Comparisons between each group were analyzed using a Student’s T-test (**** P < 0.0001, ns, P ≥ 0.05).

### Knockdown of CTU1 affects erythroid differentiation

We extracted the red erythroid population of interest for our research (Fig 6A). After performing dimensionality reduction clustering, we discovered that there are two distinct populations with erythroid differentiation function. Therefore, these two populations were analyzed in a pseudo-time series using the reference cell population. The pseudo-time series results showed that the final mature populations of the two groups were different. The mature cells in the wild-type group belonged to cluster1, while the mature cells in the mutant group belonged to cluster0. The cells in the mutant group were generally in the early stage of differentiation (Fig6B,C). In order to gain a better understanding of the role of CTU1 in erythroid cells, we conducted a differential gene analysis. The results revealed that the downregulation of CTU1 expression would result in increased DNA repair and promotion of erythroid differentiation. Additionally, it would lead to a decrease in translation function and the cell’s response to starvation (Fig 6D, E, SupplementalTable S9,S10). Erythroid cell-specific regulatory factors include LMO2, HLF, GATA1, and TAL1 (Fig SI.2E). Compared to the wild-type cells (Fig. 6F), we identified regulators with high activity, including E2F7, E2F8, HCFC1, and HCFC2, which play important roles in hematopoiesis[43–45]. RB1, TFDP1, and EZH2 affect the development of erythroid cells[46–48]. In contrast, the activity-regulating factor KLF11[49], which is expressed in mature red blood cells, and the factors SREBF2, CREB3, and ATF6 involved in endoplasmic reticulum stress[50–52], as well as the NFKB2 factor that plays a role in early differentiation of the bone marrow lineage[53], all show decreased activity. These results suggest that there are specific regulons that are active in erythroid cells, and their activity differs between wild-type (WT) and knockdown CTU1. This finding reinforces the heterogeneity of erythroid cells and highlights the potential importance of CTU1 in erythroid biology.

**Figure 6.**
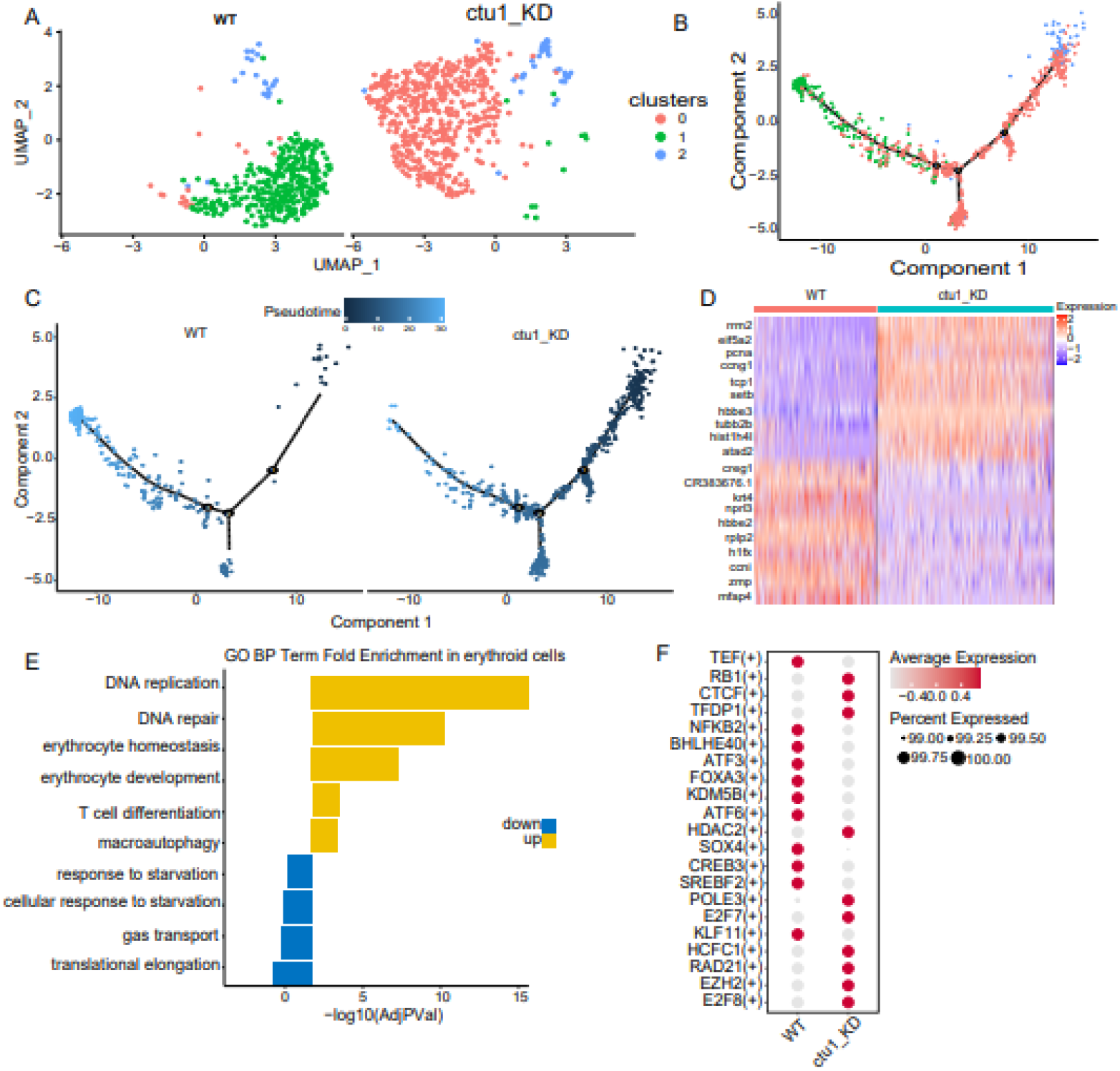
Heterogeneity of ctu1 defects in wild zebrafish erythroid cells. (A) UMAP visualization of erythroid cells, colored according to clusters. (B) Cell trajectory plot of erythroid cells. Cells are colored based on clusters. (C) Pseudotime trajectories of erythroid cells. (D) Heatmap plot of the top 10 DEGs between WT and Ctu1−/− erythroid cells. (E) The bar plot shows the GO BP analyses of DEGs between WT and ctu1−/− erythroid cells. The y-axis represents the enriched BP term, while the x-axis represents the −log10 (adjusted P-value). GO terms enriched in ctu1-/- are colored yellow, and terms enriched in WT are colored blue. (F) Dot plots show changes in the expression of transcription factors across different samples. The color and size of circles indicate the average expression level and percentage of cells.

## Discussion

In this study, we knocked down the expression of Ctu1 in zebrafish and combined it with scRNA-seq. We discovered, for the first time, that tRNA thiolase CTU1 plays a role in zebrafish embryonic development, angiogenesis, and erythroid differentiation. It primarily regulates ribosomal genes and certain adhesion factors.

tRNA is a small molecule nucleotide RNA that is abundant in cells and plays an important role in protein translation[1, 54]. It is also the nucleic acid with the highest number of modified components, with most modifications occurring at the 34th and 37th sites of the anticodon stem loop[55, 56]. These modifications can impact the tertiary structure folding, stability, and function of tRNA, which is crucial for ensuring the efficiency and accuracy of protein translation and maintaining appropriate cellular protein stability[56–58].

The Elongator complex is highly conserved in eukaryotes, and mutations in its six subunits are associated with neurological diseases[59, 60]. For example, Elp2 negatively affects the activity of the complex and its translation function through tRNA modification, leading to microcephaly and nerve damage in mice[61]. The pathogenic biallelic missense mutation in ALKBH8 is associated with an autosomal recessive form of neurological diseases syndrome[62]. A homozygous CTU2 mutation was found in patients with a novel multiple congenital anomalies syndrome called DREAM-PL, which consists of dysmorphic facies, renal agenesis, ambiguous genitalia, microcephaly, polydactyly, and lissencephaly [63, 64]. After knocking out CTU1, we observed that zebrafish had brain edema, which may be attributed to abnormal neurodevelopment. Combined with single-cell data analysis, the down-regulated genes of nerve cells were found to be enriched in the growth cone and polarized growth pathways. This suggests that the deletion of CTU1 may affect the formation and guidance of the growth cone. Affected, which in turn affects the growth and connection of neuronal axons[65]. This eventually leads to abnormal development of the nervous system, including errors in neuronal connections, errors in axon path selection, or defects in the formation of neural circuits[66].In light of the intricate nature of neural development, additional research and a greater amount of experimental evidence are still required. Together, tRNA modification regulates gene expression, which in turn regulates disease occurrence and physiological processes.

According to our scRNA-seq analysis, we identified 22 cell types, which is consistent with the results of two recently published articles[67, 68]. These cell types include typical nerve cells, epidermal cells, mesoderm cells, neural crest cells, optic nerve cells, erythroid cells, and endothelial cells. Because angiogenesis, obstruction, and cardiac hypertrophy were observed in zebrafish animal models, which involve cells belonging to the mesoderm cell subgroup, we conducted further studies on this subgroup. The mesodermal cells were selected for further study. Single-cell transcriptome data showed that the proportion of cells in the G2/M and S phases was higher after ctu1 knockdown compared to the WT group. Differential gene and GSVA analysis revealed the inhibition of angiogenesis, vascular endothelial growth factor receptor binding, and migration of endothelial cells. This is consistent with the phenotype observed in zebrafish after CTU1 deletion.

We further extracted endothelial cells for analysis. We found that knocking down ctu1 significantly inhibited the expression of ribosomal genes (such as rpl13, rpl15, and rpl18) in endothelial cells compared to the wild type. Changes in the abundance or function of specific tRNAs can lead to codon-biased reprogramming of mRNA translation, resulting in a high degree of regulation of protein synthesis, energy metabolism, and cell death. This can also lead to changes in global protein expression[57, 69]. We propose a hypothesis that tRNA thio-modifying enzymes are involved in ribosomal transport mechanisms and, as a result, impact angiogenesis. This hypothesis could be a potential avenue for future exploration. Additionally, we analyzed erythroid cells and observed that, in the early stages of cell differentiation, after CTU1 knockdown in pseudo-time series, differential gene enrichment analysis revealed that up-regulated genes were enriched in erythroid development, hemoglobin, and erythroid differentiation pathways. It is speculated that erythroid cells deficient in CTU1 are in the primitive stage of red blood cell development[70].

In addition, cell communication plays a crucial role in the process of development. Our cell communication results indicate that knockdown of ctu1 affects the expression and activity of endothelial cell adhesion factors ESAM and CDH5. Both factors are simultaneously inhibited, leading to the disruption of endothelial junction structure and potentially fatal blood clotting[71]. The signaling pathways that decrease after CTU1 deletion also include NOTCH and VEGF signals, both of which are involved in angiogenesis[72, 73]. The specificity of WNT signaling significantly increased after CTU1 deletion. During early embryonic development, the Wnt/β-catenin signaling pathway regulates the cell fate of embryonic endoderm cells and influences the development of vertebrate organs [74].

In summary, our study will further promote the functional study of the tRNA-modifying enzyme CTU1 at the mammalian level. Using zebrafish as model organisms, combined with scRNA-seq analysis, researchers initially discovered that deletion of CTU1 could result in zebrafish embryo dysplasia, blocked angiogenesis, and inhibited erythroid differentiation.

## Author contributions

YZW Yu did the single-cell data analysis. Y Wang, H Shi and CQ Wang performed the experiments. YZW Yu, CQ Wang, and ZL Zhou wrote the manuscript. ZL Zhou supervised the project.

## Supplementary information

Supplementary information is available at the website of Development.

